# Functional resurveys and models reveal the interplay of plasticity and evolution of Pierid butterflies in response to recent climate change

**DOI:** 10.1101/2025.09.23.678128

**Authors:** Lauren B. Buckley, Joel G. Kingsolver

**Affiliations:** Department of Biology, University of Washington, Seattle, WA 98195-1800, USA; Department of Biology, University of North Carolina, Chapel Hill, NC 27599, USA

**Keywords:** adaptation, butterfly, climate change, insect, plasticity, trait

## Abstract

The extent of contemporary evolution, which is mediated by interactions with plasticity, will be an important determinant of biological responses to climate change. We synthesize two functional resurvey projects that, coupled with mechanistic models, evaluate the interplay of plasticity and evolution of Pierid butterfly larval (thermal sensitivity of feeding) and adult (wing melanization) traits over recent decades. We characterize thermal environments over the resurvey periods, which we interface with developmental and (historical, current, and hypothetical) thermal sensitivity traits to examine the implications of evolutionary changes. We find that the evolution of photoperiod-cued plasticity of wing melanization in California *Colias* is consistent with avoiding thermal stress during warming springs. Plasticity has not evolved for Colorado *Colias* populations, which have experienced stronger increases in climate means relative to extremes in recent decades. Evolution in Colorado *Colias* larvae has improved tolerance to warm extremes, whereas evolution in California *Colias* larvae has broadened thermal sensitivity consistent with capitalizing on expanded seasonal thermal opportunity. Our models predict that Washington *Pieris* larvae have experienced shifts in the direction of selection to increase performance at warm temperatures. The research highlights the importance of evaluating changes in climate change exposure and sensitivity to understand interacting organismal responses.

## Introduction

The biological responses of organisms to recent and ongoing climate change are increasingly obvious (Halsch et al. 2021; Harvey et al. 2023). Population declines, changes in seasonal timing, latitudinal and elevational shifts in geographic range, and even extinction have now been documented in numerous taxa (Scheffers et al. 2016). Despite these general patterns, the responses of different populations and species are often heterogeneous and can appear unpredictable (Maguire et al. 2015; Beissinger and Riddell 2021). In addition, adaptive phenotypic plasticity and evolution can play potentially important roles in reducing the fitness consequences of climate change for some species (Chevin et al. 2010; Urban et al. 2024). Understanding and predicting the diverse responses of different populations and species to ongoing climate change remains a major challenge for biologists.

One conceptual (and mathematical) way to view this challenge is in terms of the relationship between environmental (e.g. climate) conditions and the optimal value of some phenotype that adapts the population to the environmental conditions (fig. 1). Climate change over time leads to a ‘mismatch’ or lag between climate conditions and the mean phenotype, reducing mean population fitness that can lead to population declines and (ultimately) extinction. Given phenotypic and genetic variation in the phenotype, the mismatch generates directional selection and phenotypic evolution that reduces (but does not eliminate) the lag between mean and optimal phenotype. Models show that the evolutionary capacity for the population to track the changing climate will depend on the rate of climate change, phenotypic and genetic variation, population size, and stochastic variation in climate (Lynch and Lande 1993; Bürger and Lynch 1995; Kingsolver and Huey 1998; Chevin et al. 2010).

**Figure 1.**
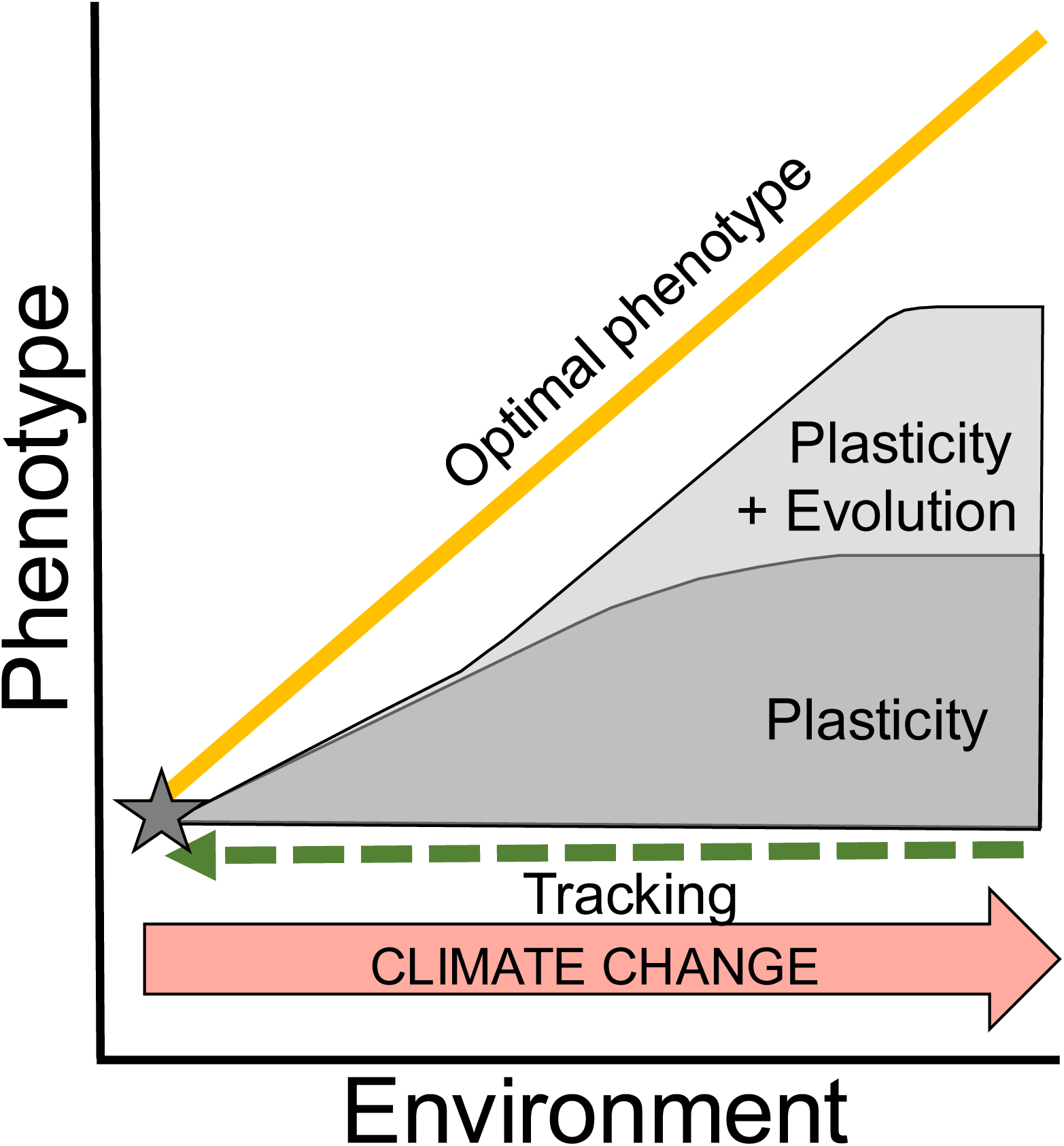
An illustration of how climate change responses interact to align phenotypes with environments. Climate change can cause a shift along the line of optimal phenotypes (solid line) such that over time, the mean population phenotype (star) become misaligned with the optimal phenotype. A common and oft studied response is environmental tracking through space or time to realign the phenotype with the optimum (dashed arrow). Phenotypic plasticity can initially allow the population to closely (but imperfectly) track optimum phenotype, but limits to plasticity will eventually to an increasing lag between the mean and optimal phenotypes. We depict a hypothetical trajectory of phenotypic plasticity as the primary process and then consider how adaptive ecology can reduce the gap to the optimal phenotype.

In this framework, adaptive plasticity plays an important role by reducing the mismatch between environmental conditions and mean phenotype (fig. 1) and reducing the fitness consequences of climate change. As a result, plasticity can ‘buy time’ for adaption by promoting persistence, maintaining genetic variation, and revealing cryptic genetic variation (Ghalambor et al. 2007; Snell-Rood et al. 2018; Fox et al. 2019). However, by reducing directional selection plasticity can delay adaptive evolution in response to climate change (Kingsolver and Huey 1998, 1998; Chevin et al. 2010, 2010). In addition, plasticity is rarely perfect and is not unlimited (Murren et al. 2015), so that the lag between optimal and mean phenotype will eventually widen with ongoing climate change (fig. 1). Finally, if adaptive plasticity itself can evolve, this can further reduce the mismatch between population and environment and maintain population persistence.

Recent studies that repeat historical lab and field experiments provide important case studies of plastic and evolutionary responses to recent climate change (Buckley and Kingsolver 2021). For example, the temperature-size rule for ectotherms—warmer developmental temperatures produce smaller adult sizes—has resulted in smaller mean size in numerous organism in response to recent climate change (Gardner et al. 2011; Sheridan and Bickford 2011). Functional resurveys with birds, plants and insects have documented evolutionary shifts in thermal sensitivity, coloration and behavior in response to climate warming during the past four decades (Higgins et al. 2014; MacLean et al. 2019; Nielsen and Kingsolver 2020; Wooliver et al. 2020). In addition, evolution changes in plasticity—specifically, in the traits that determine the seasonal timing of diapause and migration in relation to climate—have occurred in both insects and birds in response to recent phenological shifts due to climate change (mosquitoes, migratory flycatchers) (Bradshaw and Holzapfel 2007; Helm et al. 2019). These accumulating functional resurveys illustrate the importance of plasticity and evolution for adaptation to climate change in at least some species.

Despite this useful, simple framework (fig. 1), climate change is much more than increasing mean temperature: it involves changes in patterns of temperature and precipitation at multiple temporal and spatial scales (Masson-Delmotte et al. 2021). This makes it difficult to directly connect plastic or evolutionary changes in traits with particular aspects of climate change. More generally, patterns of environmental variability are central to the interplay of plasticity and evolution (Chevin et al. 2010), and the predictability of environmental cues is critical for adaptive plasticity and its evolution (Bonamour et al. 2019; Leung et al. 2020). Understanding how populations adapt to the complexities of climate change in natural environments will require both historical resurveys that document changes in phenotypes and environments, and a quantitative, mechanistic framework for connecting environment, traits, variation and selection (Kingsolver and Huey 1998; Buckley et al. 2023).

In this paper, we synthesize and extend research combining historical resurveys and mechanistic models to explore the interplay of phenotypic plasticity and evolution in response to climate change in insect populations. Insects offer an excellent system for resurveys due to their high sensitivity to environmental conditions, their amenability to lab studies, and extensive historical records. Dramatic recent declines in insect populations (Halsch et al. 2021; Wagner et al. 2021; Harvey et al. 2023), including of butterflies in the Western United States (Forister et al. 2021), highlight their sensitivity to recent environmental change. Our studies with Pierids focus on two key phenotypes— wing melanism in adults and thermal performance curves (TPCs) for larvae—that mediate fitness determining responses to thermal environments. The multiple generations per year of most pierid species experience different thermal conditions. Plasticity in development time and wing coloration can buffer variation in selection across seasons and years (Kingsolver and Buckley 2015, 2017).

In this paper we build on past resurvey results documenting evolutionary changes in the plasticity of wing coloration, in mean wing coloration, and in larval TPCs over the past 30-50 years (Higgins et al. 2014; MacLean et al. 2019; Nielsen and Kingsolver 2020). We integrate detailed climate data with microclimate models to characterize shifts in thermal environments across the resurveys. We then interface the thermal conditions with developmental traits and (historical, current, and hypothetical) TPCs to examine the implications of the evolutionary changes. We estimate shifts in mean performance and stressful extremes both seasonally and in response to recent climate change. We hypothesize that recent evolutionary changes have enabled capitalizing on expanded thermal opportunity while alleviating thermal stress.

### Study species and thermal biology

Our resurvey studies and modeling have focused on four Pierid butterfly species in North America, in the genera *Colias* (‘Sulphurs’) and *Pieris* (‘Whites’). The *Colias* resurveys focused on three species: *C. meadii*, which is restricted to high-elevation meadows (∼3200-3700m) in the Rocky Mountains (Watt et al. 1977); *C. eriphyle*, which is present at lower elevations (1400-3000m) in Colorado (Watt et al. 1979); and *C. eurytheme*, which is present at low elevations including in California and is closely related to *C. eriphyle*. *C. meadii* is univoltine (a single generation per year) with an obligate larval diapause, while *C. eriphyle* and *C. eurytheme* are multivoltine (multiple generations per year) in the study areas (Ae 1957, 1958; Stanton 1982). *Pieris rapae* L. is native to Europe and is now widespread across much of North America, where it is generally multivoltine (Ryan et al. 2019).

The larval and adult stages of these Pierids differ importantly in their thermal biology and adaptations to local climate. The Pierid larvae do not actively regulate body temperature except at extreme high temperatures and do not exhibit variation in larval coloration relevant to thermal biology (Sherman and Watt 1973; Kingsolver 2000; Kingsolver et al. 2004; Higgins et al. 2014).

This contrasts strongly with the thermal biology of adult Pierids. Flight in Pierid butterflies is restricted to a relatively narrow range of body temperatures, which often exceeds environmental temperatures in their montane environments (Kingsolver 1983, 1985). Flight is essential to fitness determining processes, including foraging, mating, and reproduction. Flight time closely indexes reproductive output because Pierids lay eggs singly on host plants (Kingsolver 1983). A single adult phenotype—wing coloration—shapes butterfly responses to thermal environmental variation at both acute and chronic timescales. By using thermoregulatory posture to alter exposure of dark wing regions to radiation, Pierids can behaviorally adjust body temperatures and thus flight duration (Kingsolver 1985). In response to thermal extremes, dark wings can cause overheating and damage, including sharp declines in egg viability (Kingsolver and Watt 1984). In Pierid species with multiple generations per year, melanism is seasonally plastic, with (for example) darker wings in cool seasons (short photoperiods, cooler developmental temperatures) (Watt 1969; Shapiro 1976). Such seasonal plasticity enables adults to balance flight capacity against risk of overheating in different seasonal weather conditions (Kingsolver and Wiernasz 1991).

### Climate data and microclimate model

We analyze thermal shifts over time for three locations relevant to understanding the evolution of larval and adult traits in response to recent climate change: Sacramento Valley, CA (38.44N, 121.86W); Montrose County, CO (38.62N, 108.02W); and Seattle, WA (47.65N, 122.29W). Environmental conditions can vary considerably between weather station and plant height and between sun and shade (Buckley et al. 2018; Pincebourde and Casas 2019; Briscoe et al. 2023). Here we characterize conditions at plant height using the request_era5 and micro_era5 functions in the NicheMapR R package (Kearney and Porter 2017), which utilize the mcera5 package (Klinges et al. 2022) to download hourly environmental data and scale them to the height of larvae on plants using a microclimate model. The ERA5 data are generated across a 0.25° latitude x 0.25° longitude grid by assimilating observational weather data into a forecast model and accessed via the Copernicus Climate Change Service (https://climate.copernicus.eu/climate-reanalysis). The data afford reasonable spatial resolution, high temporal resolution, and a complete time series for temperature, solar radiation, and windspeed. We assumed the insects perched on vegetation at heights of 0.2m. We additionally assumed ground reflectance of 30% (REFL=0.3) and a surface roughness height of 0.02m (RUF=0.02), corresponding to habitat measurements for Colorado *Colias* (Buckley and Kingsolver 2019). We estimated radiation using the ERA5 data (IR=2), but we checked that estimated temperatures were similar using the NicheMapR algorithms. We ran the microclimate model to estimate air and ground temperatures at plant heights assuming 25% shade (minshade=25). Figure S1 illustrates the temperature differences between reference and plant height for our focal sites.

### Adult wing melanization

#### Evolution of wing melanization

We synthesize past findings from developing and testing a mechanistic modeling framework for *Colias* that incorporates microclimate, heat balance, and demographic models (Buckley and Kingsolver 2012) as well as evolution (Kingsolver and Buckley 2015) and plasticity (Kingsolver and Buckley 2017) to project selection on adult wing traits through time. The model projected that evolutionary selection favors wing lightening at low elevation (to avoid overheating) but wing darkening at high elevation (to capitalize on thermal opportunity) across the butterflies’ distribution (Kingsolver and Buckley 2015). Importantly, however, we found that seasonal and annual variation in climate causes the strength and direction of selection to fluctuate (Kingsolver and Buckley 2015). Our models suggest that plasticity in wing absorptivity can facilitate evolution, particularly at lower elevations with long seasons, by reducing temporal variation in the strength and direction of evolutionary selection (Kingsolver and Buckley 2017). Phenological shifts (e.g., timing of maturation) caused by environmental effects on developmental rate can also reduce variation in selection (Kingsolver and Buckley 2018).

Projections of responses to future climate change indicate that some regions will experience shifts in the direction of selection over time; initial selection for wing darkening to extend daily and seasonal activity periods and capitalize on warming in cool areas will eventually—as climate change proceeds —shift to selection for wing lightening to avoid overheating (Buckley and Kingsolver 2019). Projected evolutionary lags may reduce fitness, but we projected that plasticity and the evolution of plasticity will shorten these lags (Buckley and Kingsolver 2019). Testing with lab and field experiments (Higgins et al. 2014, 2015; MacLean et al. 2016a, 2016b) and museum specimens (MacLean et al. 2016c, 2019) partly confirmed model predictions but also highlighted how the interactions of plasticity and evolution complicate phenotypic shifts (Buckley and Kingsolver 2019). Shifts in wing melanism among museum specimens of *Colias* over the past half century generally supported model predictions (MacLean et al. 2016c, 2019).

#### Evolution of plasticity in wing melanization

We examine the differential evolution of plasticity in wing melanization between the related Colorado *C. eriphlye* and California *C. eurytheme*. Contrary to model predictions for Colorado *C. eriphyle* (Kingsolver and Buckley 2017), we detected no evolution of plasticity in wing absorptivity, perhaps because genetic constraints were omitted from the model (MacLean 2015). We assess how the evolution of reduced melanism in California *C. eurytheme* at short photoperiods over nearly 50 years corresponds to thermal changes (Nielsen and Kingsolver 2020). Warming has been concentrated in spring and modest in the summer (fig. 2A).

**Figure 2.**
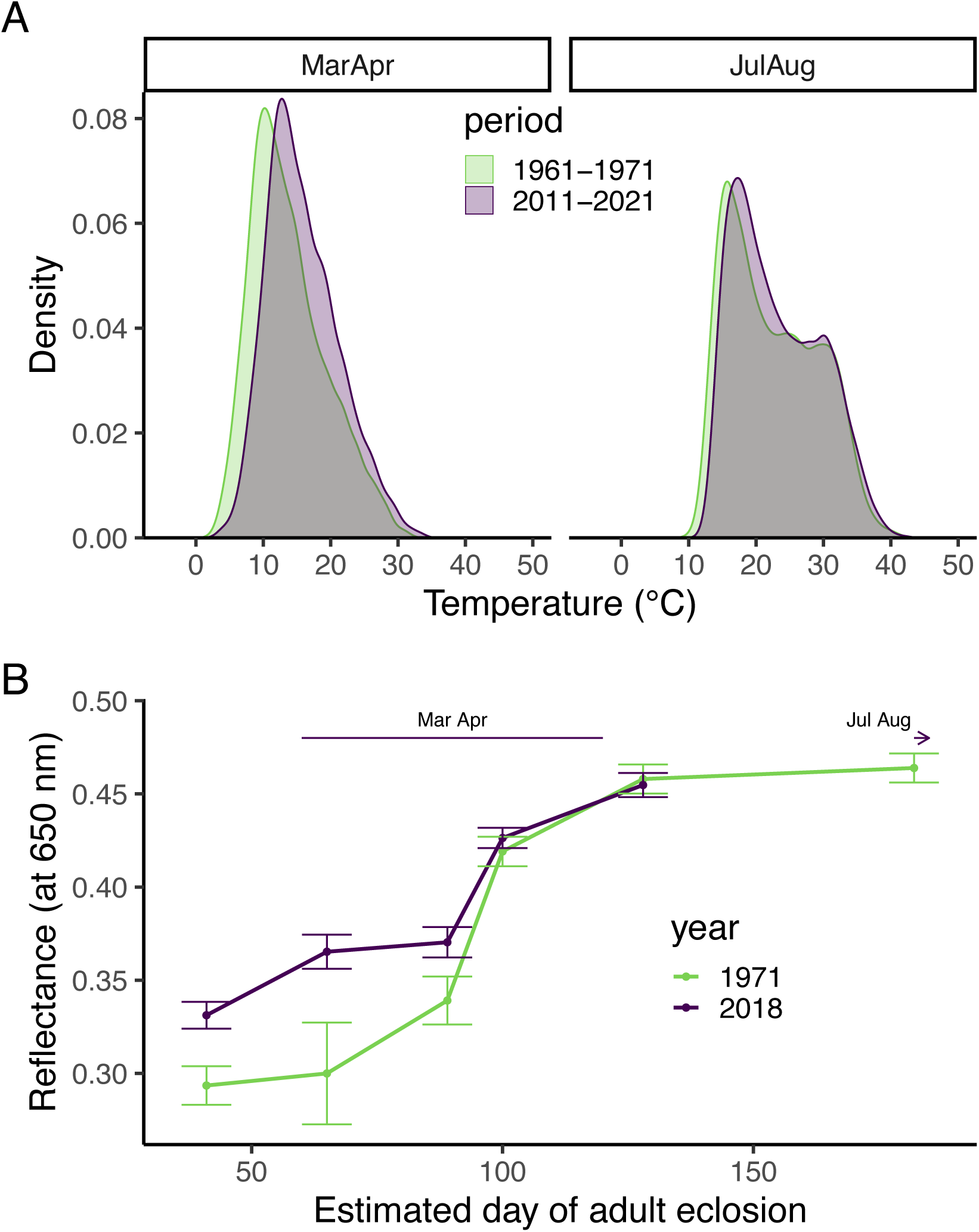
(A) Daytime hourly temperatures at plant height have increased over recent years (2011-2021) for California *Colias eurytheme*. Warming has been concentrated in the spring (March to April). (B) Ventral hindwing wing reflectance (means ± 95% CI) of male *C. eurytheme* has increased at short photoperiods between 1971 and 2018 (Nielsen and Kingsolver 2020), consistent with selection for lightening adults of the seasonal generation that have experienced the most pronounced warming.

We estimated the day of adult eclosion corresponding to the photoperiods used in the laboratory study. Laboratory data for *C. eurytheme* at constant temperatures from 15 to 35°C at 5°C increments (Allen and Smith 1958) enabled quantifying pupal development time. We estimated T_0_, the lower development temperature (°C), and DD, the number of degree days to complete the development, as -*b*/*a* and 1/*a*, respectively, where *a* is the slope and *b* is the y-intercept of a linear regression with temperatures along the x-axis and development rate (1/duration of development) on the y-axis (Trudgill et al. 2005). We calculated the duration of pupal development using temperatures at plant height in the shade to estimate the number of days until accumulated growing degree > DD. Roughly corresponding to experimental data, we assumed a maximum duration of pupal development range of 40 days and the shortest estimated duration was 9 days at the long photoperiod.

Plasticity has evolved such that wing reflectance has increased (absorptivity and melanization has decreased) for the first season generation (historically mid-April) but has not changed for subsequent generations (fig. 2B). Historically the second generation occurred in June and the third in July and August (Allen and Smith 1958). Thus, the evolution of plasticity is consistent with decreasing body temperatures (via increased reflectance) to avoid thermal stress in response to increasing spring temperatures, while maintaining similar body temperatures in summer, when temperatures have remained relatively stable (Nielsen and Kingsolver 2020).

High levels of gene flow may have prevented the evolution of plasticity for Colorado C. *eriphlye* (MacLean 2015), but another factor may have been the greater reliance on temperature rather than photoperiod relative to California C. *eurytheme.* Photoperiod is a reliable cue for the thermal conditions adults experience over the long active season in California. Temperatures are a less reliable cue (Kingsolver and Huey 1998) for selection associated with thermal extremes in Colorado, which have been increasing over recent decades.

### Larval thermal sensitivity

#### Thermal performance curves and fitness outcomes

We fit thermal performance curves (TPCs, fig. S2) to short term growth or feeding rate data for *C. eriphyle* and *C. euytheme* (Higgins et al. 2014) and *P. rapae* (Kingsolver 2000) to estimate the implications of TPC evolution in response to climate change. We selected a beta formulation of TPCs (Asbury and Angilletta 2010) because it includes interpretable parameters for position (T_min_, the lower thermal limit for performance) and breadth (the range of temperatures over which performance is possible). It also enables changing TPC height along with breadth such that the area under the curve remains fixed (specialist-generalist tradeoff). We modeled performance, Z, as a non-linear function of body temperature in Kelvin, T_b_ (Asbury and Angilletta 2010):

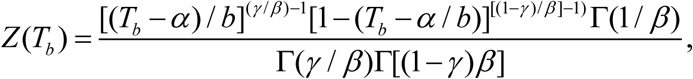

where α, β, and γ determine the T_min_, breadth, and skewness of the performance curve, respectively (fig. S2). The parameter b determines the tolerance range (T_max_ – T_min_, where T_max_ is the upper thermal limit for performance). The influence of parameter values is depicted in Figure S2. We used the nls function in R (stats package) to fit the TPCs to performance (feeding or growth rate) data. We used the rTPC R package (Padfield and O’Sullivan 2022) to confirm that the formulation fits well relative to other TPC shapes including Gaussian and Weibull.

We generated a series of hypothetical TPCs varying the position (T_min_) of the TPC from fifteen degrees below to ten degrees above the fitted values (Table 1). We additionally considered three breadths, an index of TPC curvature (fig. S2): the fitted value as well as 0.03 smaller and larger. We used the fitted values for the tolerance and skew. We assessed performance for the series of hourly body temperatures and each hypothetical TPC spanning from the historic to the recent studies. We approximated larval body temperatures as air temperatures in 25% shade at plant height, which approximates larvae occupying the undersides of the leaves (but see Pincebourde and Casas 2019). We examined what TPC thermal optima produced the highest growth rate each year. We note that our analysis does not account for the potentially detrimental effects of thermal extremes exceeding the TPC thermal tolerance. We used analysis of variance to assess the significance of trends over time and with TPC shifts.

**Table 1.**
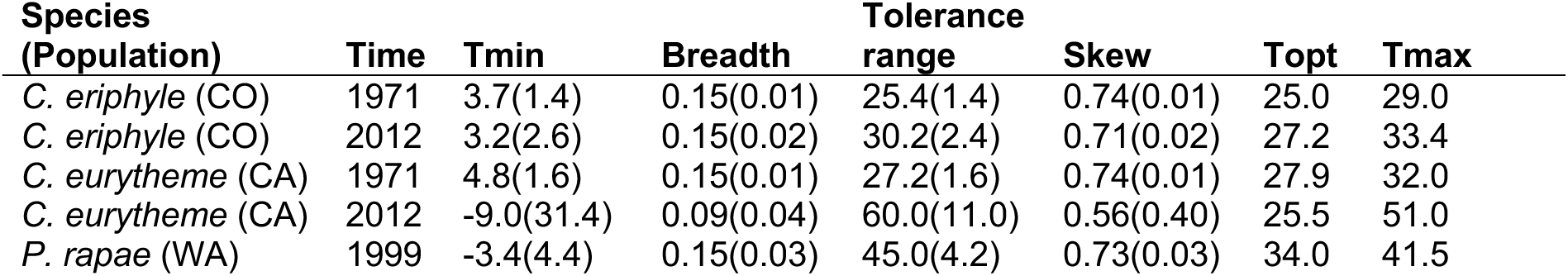
Parameter estimates and standard errors for beta thermal performance curve fits to data for the historic and recent *Colias* and *Pier*i*s* populations. We also provide estimates of the thermal optima (Topt) and upper endpoint (Tmax) of the TPC based on the fit of the other parameters.

### Colias thermal sensitivity

Temperatures at plant height have shifted over recent decades in Colorado to peak at warmer temperatures and to include a higher incidence of hot temperatures (fig. 3A). The temperature shifts correspond to an evolutionary increase in the feeding performance at warm temperature between initial experiments in 1972 and repeating the experiment in 2011 (fig. 3A) (Higgins et al. 2014). We estimate that during the initial survey *C. eriphyle* larvae would have had to use heat avoidance to avoid stressfully hot temperatures, particularly in the summer (fig. 3A). We estimate that 18% of spring temperatures and 46% of summer temperatures exceeded T_max_ in 1972.

**Figure 3.**
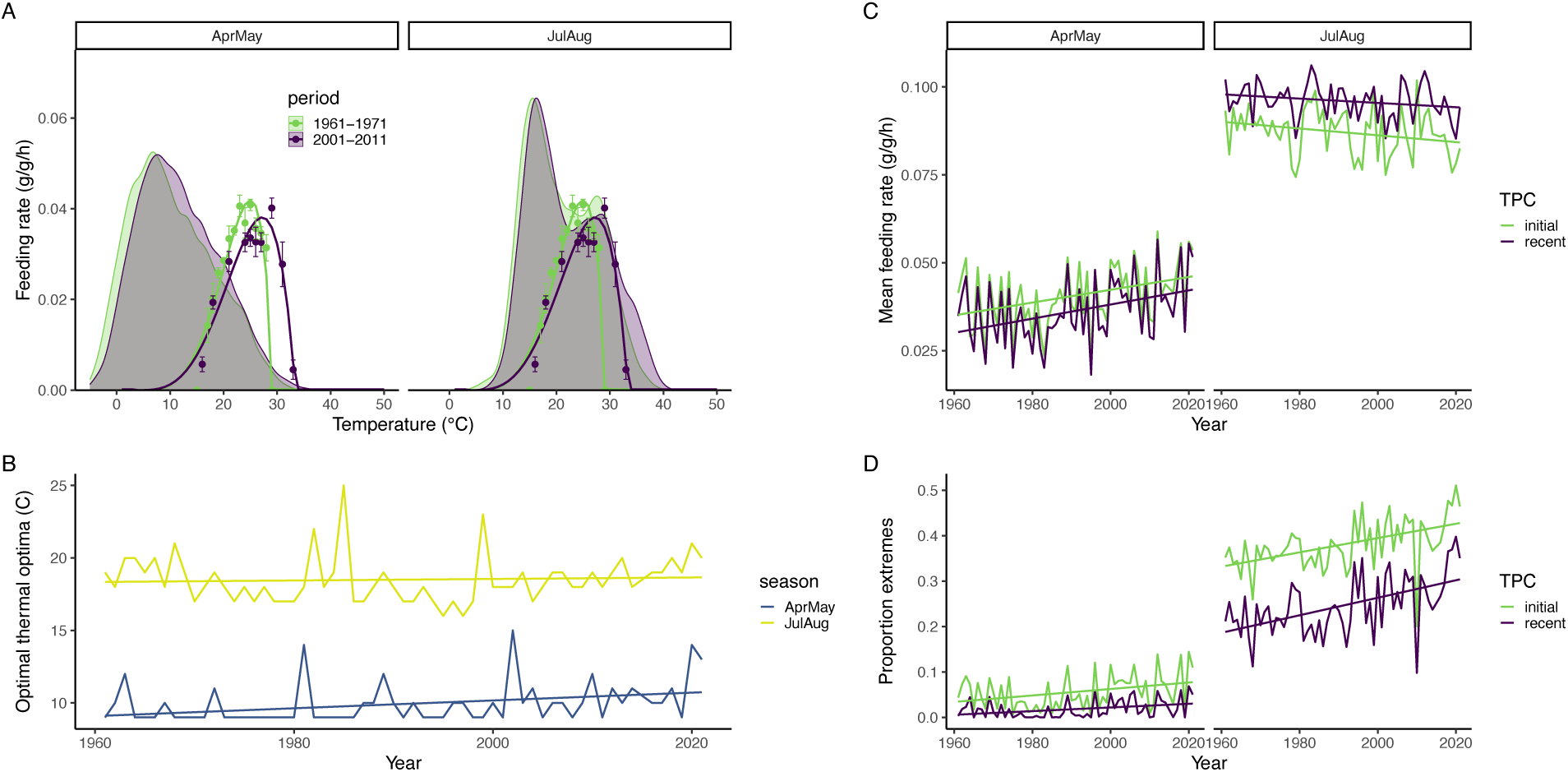
(A) The warmest temperatures at plant height have increased incidence since the initial survey, particularly during July and August for *C. eripyle* in Colorado. Overlaying the TPCs for feeding (means ± SE) aids in interpreting TPC shifts between the initial (1972) and recent (2012) experiments. (B) The thermal optima corresponding to maximal feeding is high throughout in summer but has been increasing in spring in response to recent warming. Seasonal (C) feeding rates and (D) proportion of extremes (temperatures within or beyond the upper 80% of the TPC tolerance range) vary over time differentially for the initial and recent TPC.

We apply the initial (1972) and recent (2011) TPCs to estimate performance given temperatures over time. We estimate that thermal shifts over time have increased spring feeding but decreased summer feeding (fig. 3C). Estimated decreases in the spring feeding rate associated with the evolution to the TPC observed during the resurvey (year t= 4.35, P<0.001; time period t=-2.84, P<0.01; F=13.52, P<10^−5^) have been countered by increases in summer feeding (year t= -2.62, P<0.01; time period t=8.45, P<0.001; F=39.12, P<10^−13^). We estimate that summer thermal extremes (temperatures falling in the top 20% of the TPC tolerance range) have increased substantially over time, but that TPC evolution has reduced the incidence of extremes (fig. 3D; year t= 6.53, P<0.001; time period t=-14.37, P<0.001; F=124.6, P<10^−15^). The TPC thermal optima corresponding to the highest summer feeding rate has remained relatively constant throughout the analysis period (fig. 3B; F=0.187, P=0.67). TPC breadth does not strongly influence optimal thermal sensitivity (fig. S3). For the spring period, we estimate a shift through time toward warmer thermal optima increasing feeding rate (fig. 3B; F= 7.63, P<0.01). The estimated modest shifts in mean feeding rate and thermal optima and more pronounced trends in extremes suggest selection may have been in response to thermal extremes rather than means.

For California *C. eurytheme*, the temperature distributions have remained relatively stable between the periods corresponding to the initial (1961-1971) and subsequent (2001-2011) feeding rate estimates (fig. 4A). There have been greater shifts in mean temperatures than extremes relative to the Colorado population. The TPC has broadened considerably between the experiments in 1972 and 2011 (fig. 4A) (Higgins et al. 2014). The shape of the TPC associated with broadening is not well characterized by the data as evidenced by uncertain parameter estimates (Table 1). We estimate that thermal shifts over time have increased spring feeding and summer feeding to a lesser extent (fig. 4C). TPC evolution has increased the estimated spring feeding rate (year t= 4.61, P<0.001; time period t=3.64, P<0.001; F=17.26, P<10^−6^) but decreased that in summer (fig. 4C; year t= 5.40, P=0.001; time period t=-8.46, P<0.001; F=50.33, P<10^−15^). The TPC shift has also reduced the incidence of springs with low estimated feeding rates. We estimate that the proportion of extremes has only increased slightly over time, but that the TPC shift has largely eliminated caterpillars experiencing extremes relative to their TPCs (fig. 4D; spring: year t= 4.61, P<0.001; time period t=3.64, P<0.001; F=17.26, P<10^−6^; summer: year t= 1.33, P=0.18; time period t=-91.27, P<0.001; F=4166, P<10^−15^). We estimate that the thermal optima that maximizes feeding rate has increased over time in both seasons (fig. 4B; spring: F=4.24, P<0.001; summer: F=22.34, P<0.001). The effect of TPC breadth does not shift over time, suggesting that our model does not capture the selection dynamics (e.g., thermal stress and carryover effects) that resulted in TPC broadening over recent decades (fig. S4).

**Figure 4.**
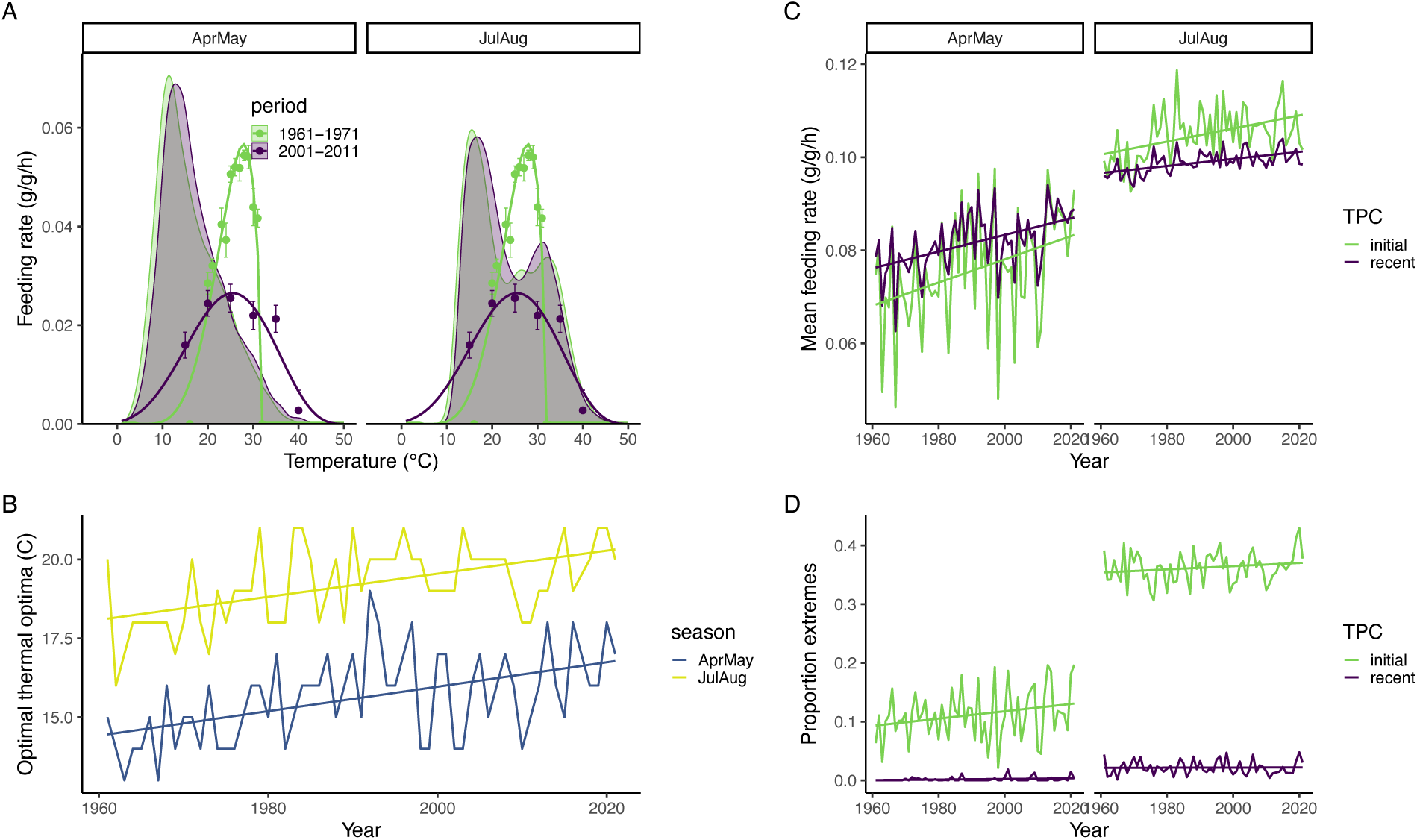
(A) The California *C. eurytheme* population has experienced more warming of thermal means than extremes at plant height relative to the Colorado population (fig. 3). We overlay the TPCs for feeding (means ± SE) to aid in interpreting TPC shifts between the initial (1972) and recent (2012) experiments. (B) The thermal optima corresponding to maximal feeding has been increasing in both spring and summer in response to recent warming. Seasonal (C) feeding rates and (D) proportion of extremes (temperatures within or beyond the upper 80% of the TPC tolerance range) vary over time differentially for the initial and recent TPC.

The *Colias* larval research highlights the insight gained from repeating historical experiments and the importance of considering shifts in climate extremes in addition to shifts in climate means. The two *Colias* populations (Colorado C. *eriphyle* and California *C. eurytheme)* exhibited differential larval responses to the increase in warm temperatures (TPC shift and TPC broadening, respectively). The differential responses may be due to the California population experiencing a longer, and thus more seasonally variable, active season and the ability to feed during both day and night (Higgins et al. 2014). Populations also differ in their ability to tolerate hot temperatures. Heat shocks differentially impact growth and gene expression in larvae from elevationally distinct populations (Higgins et al. 2015).

### Pieris thermal sensitivity

We revisit a study of selection in 1999 on the temperature dependence of feeding rate for Seattle *P. rapae* caterpillars. In that study, a selection analysis examined how larval growth rate at a series of constant temperatures corresponded to pupal mass following transplantation to the field (Kingsolver et al. 2001). The initial field study spanned July 28 – August 5, 1999 (Kingsolver 2000), and we examined data for a slightly extended period of July 26 to August 8^th^ each year. The distribution of daytime hourly larval body temperatures during the study period has shifted since the initial study to include more warm temperatures (fig. 5A). The percent of temperatures exceeding 20°C has increased from 12.3% to 13.9%.

**Figure 5.**
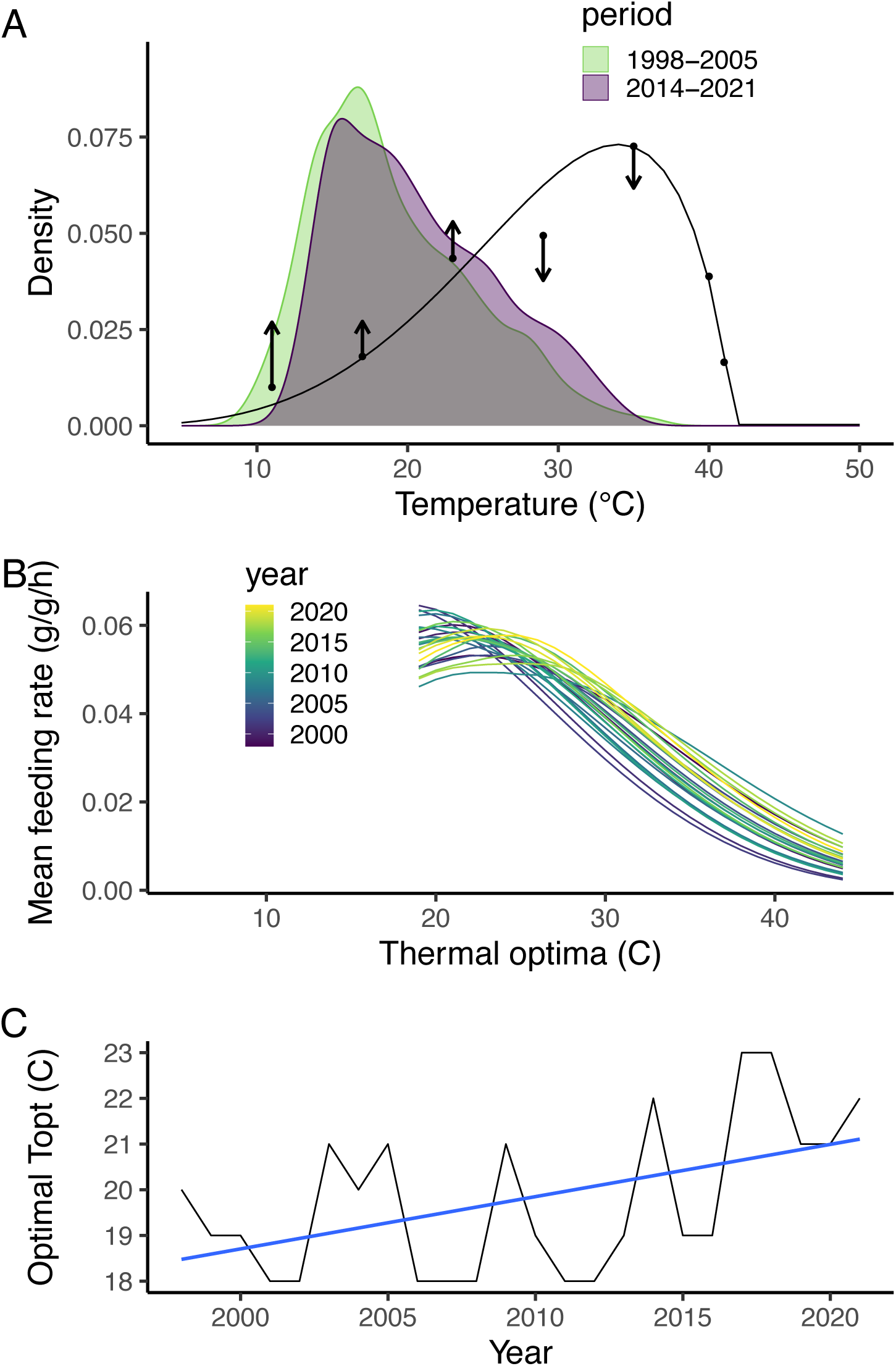
(A) Hourly larval body temperatures (July 26-Aug 8) have shifted over time to include more warm temperatures. In the 1999 study, selection acted on the thermal performance curve (TPC) for *P. rapae* larval growth (black line) to increase growth at cool temperatures and decrease growth at warm temperatures (arrow length and direction indicate selection gradients). (B) The TPC thermal optima that maximizes growth has increased over time and growth has declined overall. (C) There is considerable variability among years in the thermal optima that maximizes growth but a trend favoring warming temperatures through time.

The relatively mild summer temperatures typical of Seattle in the 1990s resulted in selection on *P. rapae* larvae that favored higher growth rates at lower (but not at higher) temperatures (Kingsolver et al. 2001) (fig. 5A). Larvae with growth TPCs such that they grew faster at cool temperatures and grew slower at warmer temperatures exhibited the largest pupal masses in the experimental gardens. We estimate that temperature shifts through time have tended to decrease *P. rapae* growth rate and shift the thermal optima associated with the highest growth rates to warmer temperatures (fig. 5B). Despite substantial interannual variability, the thermal optima favoring increase growth shifts to warmer temperatures through time (fig. 5C; F=6.76; P<0.05; see fig. S5 for the effect of breadth). These projections suggest that the selection to decrease performance at warm temperatures has decreased or reversed with climate warming. Past selection to increase growth at cool temperatures may have persisted if development initiates earlier. While the initial studies only found significant selection via pupal mass (Kingsolver et al. 2001), we anticipate that the increase in warm temperatures may have resulted in significant selection via larval survival.

We are currently repeating the initial studies to test these expectations. A key question is whether shifts in selection have resulted in TPC evolution. About ¼ of total variation in growth rates in the initial studies was attributed to genetic variation with strong genetic correlations at high temperatures, which may indicate evolutionary constraints that could limit evolution (Kingsolver 2000). We coarsely accounted for these constraints in our analysis by assuming a TPC with a fixed shape and varying thermal optima, but ongoing work is empirically evaluating how genetic correlations and variation have constrained evolution and how the constraints have shifted over time.

## Discussion

### Interplay of plasticity and evolution

The research on both *Colias* and *Pieris* butterflies reveals that environmental variability drives fluctuations in selection that can substantially slow evolution in response to climate change (Kingsolver and Buckley 2015). Even minor shifts in temperature distributions can substantially influence selection, particularly if the shifts increase the incidence of high temperatures that dramatically reduce performance or cause stress. Prominent fluctuations in selection lead to an important role for plasticity. A common assumption is that pronounced diurnal and seasonal environmental variation at high elevation will make developmental and reversible plasticity more important there than at lower elevations (Trunschke and Stöcklin 2017). The *Colias* research suggests that, while environmental variation at high altitudes can lead to a more substantial role of plasticity than at mid elevations, plasticity can play an even greater role at low elevations where extended seasons can expose generations to substantially different environmental conditions (Kingsolver and Buckley 2017). However, limited seasonal windows for the completion of development and other processes can promote plasticity at high elevations (Smith et al. 2021).

### Temporal shifts in thermal opportunity

The research also indicates the importance of considering how climate change can transiently alter thermal opportunity. Simple expectations that selection will act to enhance performance or avoid overheating at high temperatures can fail if organisms alter temporal patterns of activity (Kingsolver and Buckley 2018). The *Colias* model suggests that the fitness advantages of increased performance in formerly thermally marginal times of days or seasons can overwhelm the fitness detriments associated with overheating (Buckley and Kingsolver 2019). Resurvey expectations depend on whether seasons shift. The potential for seasonal shifts in both species will depend on the role of temperature and photoperiod in cueing diapause emergence and determining developmental rates. Another important consideration is how seasonal shifts can alter the environmental exposure of sensitive life stages (Kingsolver and Buckley 2020). Temporal shifts, particularly expanded seasons, can increase the importance of plasticity and its interplay with evolution.

Temporal shifts in thermal opportunity can lead to reversals in the direction of selection with warming. Much discussion of evolution in the context of climate change concerns whether evolution can act fast enough to rescue populations from climate change (Hoffmann and Sgrò 2011). However, the temporal shifts in the direction of selection we project for montane butterflies complicate the notion of evolutionary lags (Buckley and Kingsolver 2019). Future fitness may be reduced by past evolution if the direction of selection reverses, but plasticity may reduce this effect by shortening evolutionary lags (Buckley and Kingsolver 2019).

### Evolutionary constraints

Evolutionary constraints associated with limited genetic variation and genetic correlations can slow responses to selection (Urban et al. 2024). The *Colias* resurvey detected modest evolution of wing traits (MacLean et al. 2019) and no evolution of the reaction norm for corresponding wing traits (Higgins 2014). Data limitations prevented assessing evolutionary constraints for *Colias*, but the *Pieris* resurvey is designed to assess how genetic variation and correlations have constrained evolution. Limited data on the genetic basis of TPCs suggests that tolerance limits are moderately heritable and have considerable evolutionary potential (Logan and Cox 2020). In contrast, thermal optima and other characteristics of the middle of TPCs has low evolutionary potential (Logan and Cox 2020; Buckley and Kingsolver 2021). Genetic correlations have limited adaptation to both increases in mean conditions and variability (Logan and Cox 2020).

### Advancing understanding of the interplay of plasticity and evolution

Increases in environmental variability and season length associated with climate change (Thornton et al. 2014) are likely to select for increased plasticity, as projected for *Colias* (Kingsolver and Buckley 2017). Plasticity may be particularly important to the climate change responses of insects and other taxa with pronounced environmental sensitivity and short lifecycles (Sgro et al. 2016). However, one synthesis found no clear empirical support that plasticity in response to temperature is under selection (Arnold et al. 2019). Studies of the evolution of plasticity and its interplay with trait evolution are sparse and we have little basis for projections. Repeating historical studies of plasticity provides an expedient and promising avenue for assessing whether plasticity will evolve in response to climate change (Nielsen and Kingsolver 2020; Nielsen et al. 2023).

Phenotypic selection experiments that address tradeoffs between trait values and plasticity are also needed. Although plasticity does often trade off with thermal tolerance (Barley et al. 2021), plasticity was not altered by selection for cold tolerance in the butterfly *Bicyclus anynana* (Franke et al. 2012). Further addressing whether temperature or photoperiod predominates in cueing plasticity will aid in anticipating climate change responses (Stoehr and Wojan 2016). A productive, initial approach is to assess the reliability of cues and potential for mismatches in the context of climate change (Kingsolver and Huey 1998).

Perhaps the most important step in resolving the interplay of plasticity and evolution in response to climate change is for research to incorporate more realistic environmental variability (Buckley et al. 2023). Further incorporation of environmental variability is needed in theory as well as laboratory and field experiments (Buckley and Kingsolver 2021). Considerations of thermal history will also be important (Williams et al. 2016). For example, experiments with the seed beetle *Callosobruchus maculatus* found that the potential for plastic and evolutionary responses to environmental change depended on whether the population has experienced past selection in fluctuating environments (Hallsson and Björklund 2012). Responses of montane butterflies to recent climate change highlights that plasticity will play an important role in buffering fluctuations in selection in responses to environmental variability. The interactions of organismal processes are likely to shape responses to climate change.

## Acknowledgments

We thank the many researchers and students who contributed to the studies synthesized here, particularly Taylor Hatcher, Jessica Higgins, Heidi MacLean, Matthew Nielsen, J. Gwen Shlichta, and Adam Steinbrenner. This research was supported by the US National Science Foundation (DEB-1120062 to L.B.B. and J.G.K., IOS-2222089 to L.B.B., IOS-2222090 to J.G.K).

## Statement of Authorship

L.B.B. and J.G.K. conceptualized the research and wrote the paper. L.B.B. conducted the analyses and created the figures with input from J.G.K.

## Data and Code Availability

Data and code are available in GitHub (https://github.com/lbuckley/WApierids) and archived in Zenodo (Buckley and Kingsolver 2025).

