## Supplementary material for "Functional resurveys and models reveal the interplay of plasticity and evolution of Pierid butterflies in response to recent climate change": Figure S1

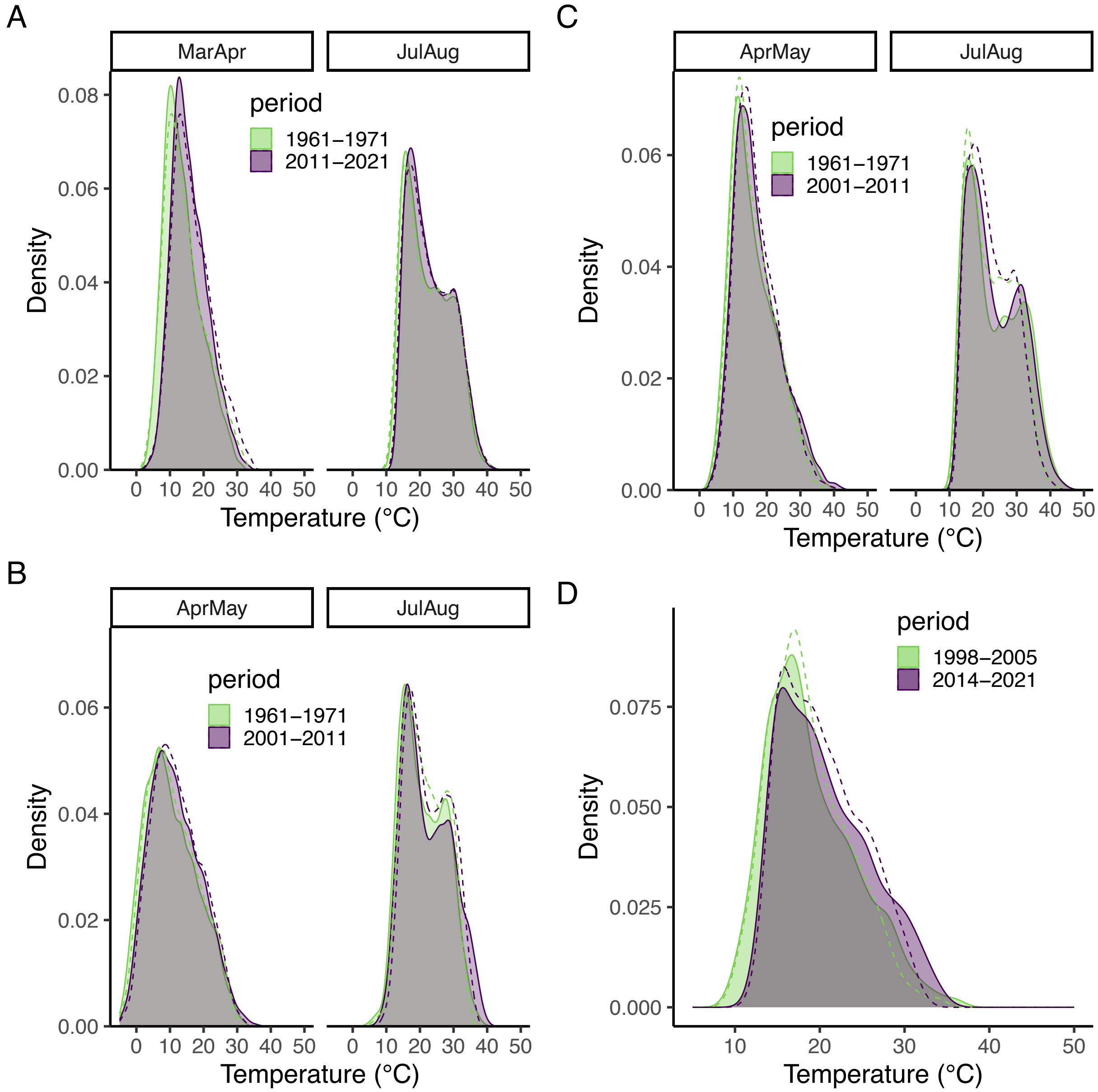


**Figure S1**. A comparison of the temperature distributions at plant (solid lines and shading) and at reference height (dashed lines) for the (A) California *Colias* resurvey of plasticity in wing melanism, (B) Colorado *Colias* resurvey of larval feeding, (C) California *Colias* resurvey of larval feeding, and (D) Washington *Pieris* resurvey.

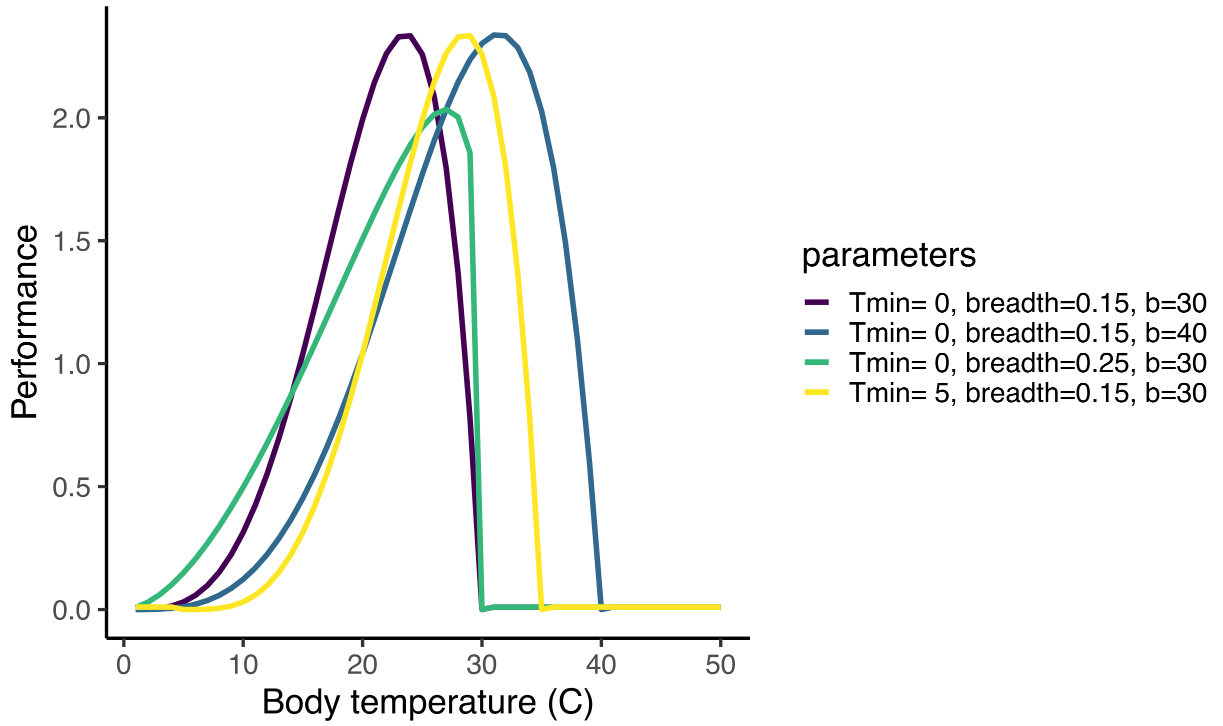


**Figure S2**. The influence of parameters (α: Tmin, β:breadth, and b: tolerance range) on TPC shape (γ:skew=0.7).

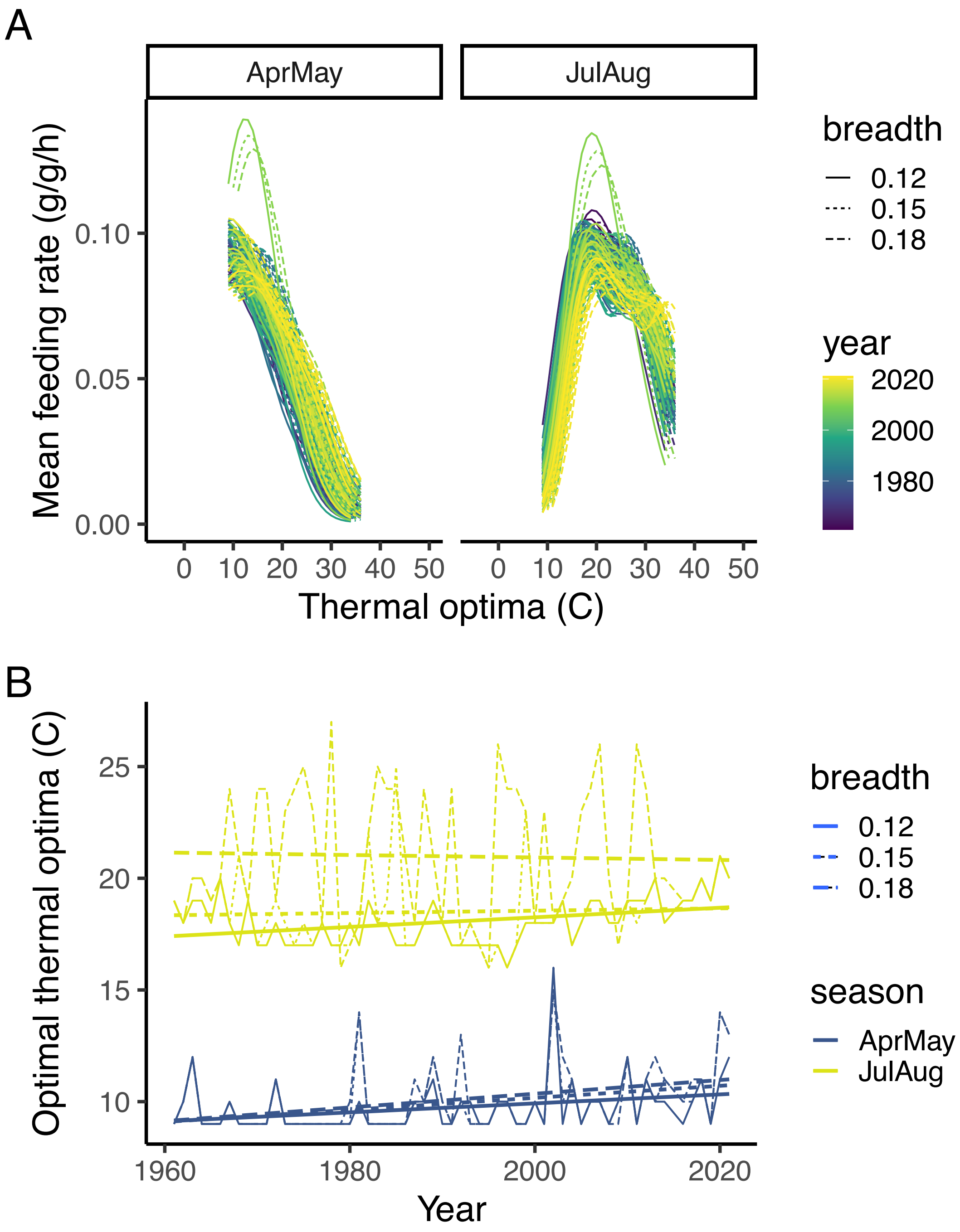


**Figure S3**. (A) We use hourly larval temperatures for Colorado *Colias* for each year to estimate how thermal optima influences feeding rate and estimate the (B) thermal optima associated with maximum feeding rate for each season. The TPC breath parameter slightly influences (A) the thermal optimal corresponding to maximum feeding and (B) the thermal optima across time.

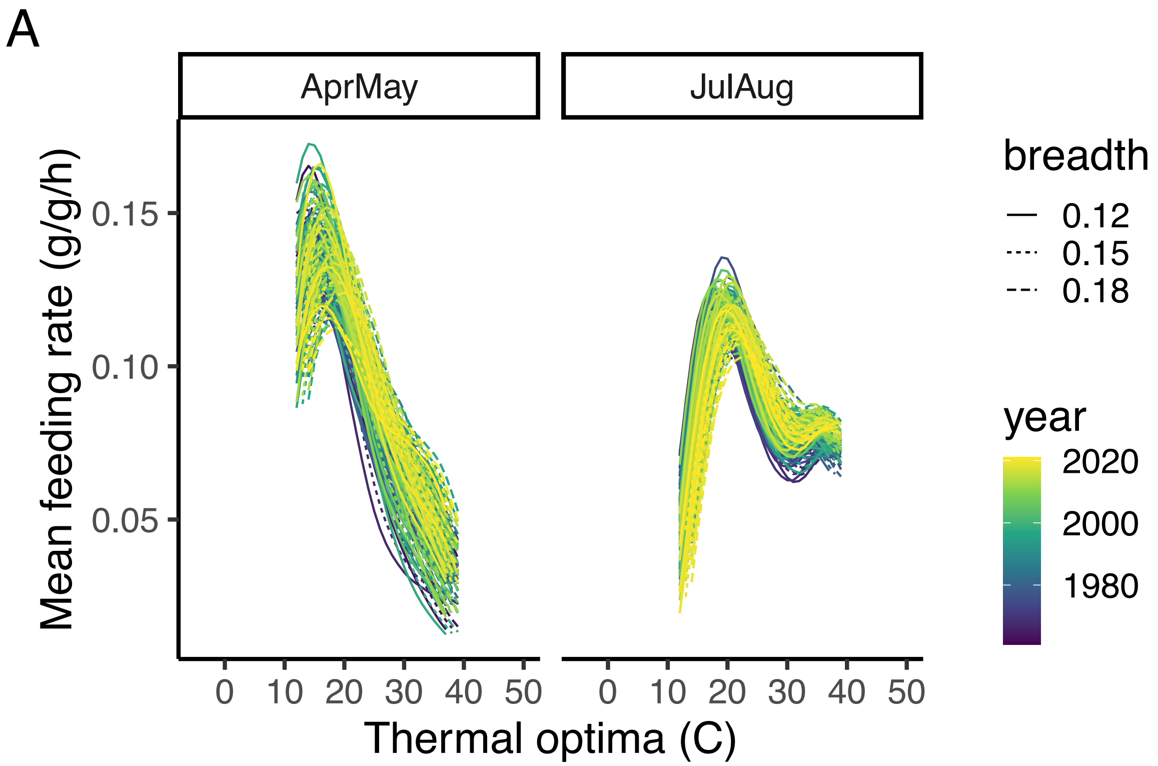


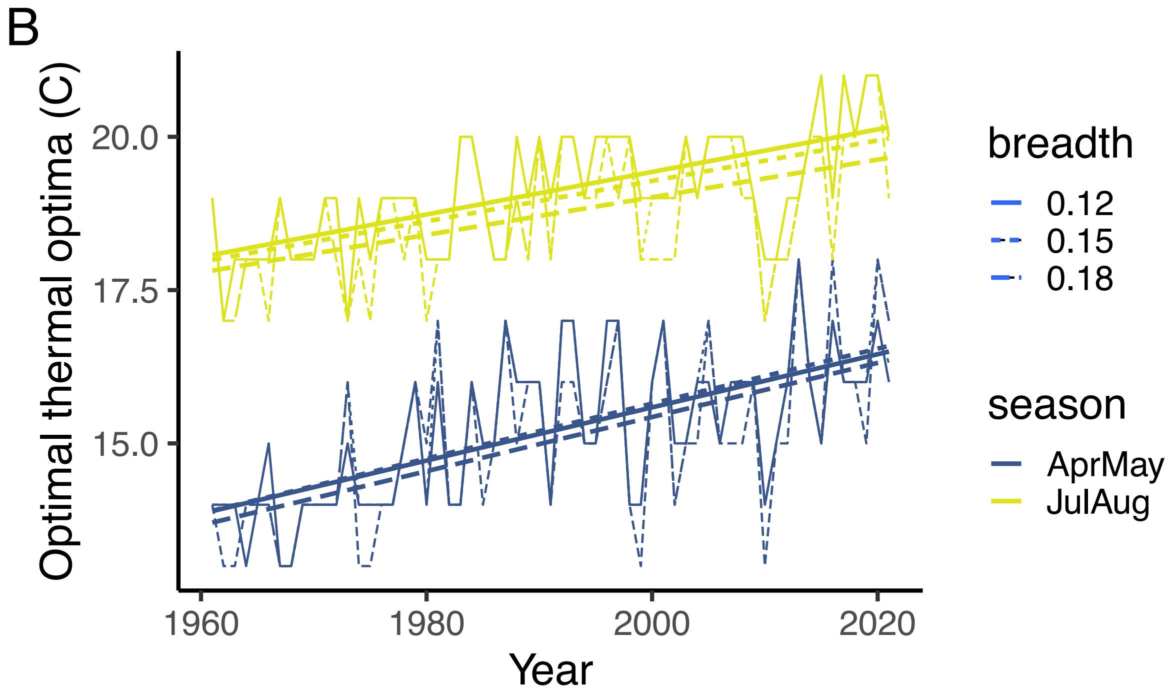


**Figure S4**. (A) We use hourly larval temperatures for California *Colias* for each year to estimate how thermal optima influences feeding rate and estimate the (B) thermal optima associated with maximum feeding rate for each season. The TPC breath parameter slightly influences (A) the thermal optimal corresponding to maximum feeding and (B) the thermal optima across time.

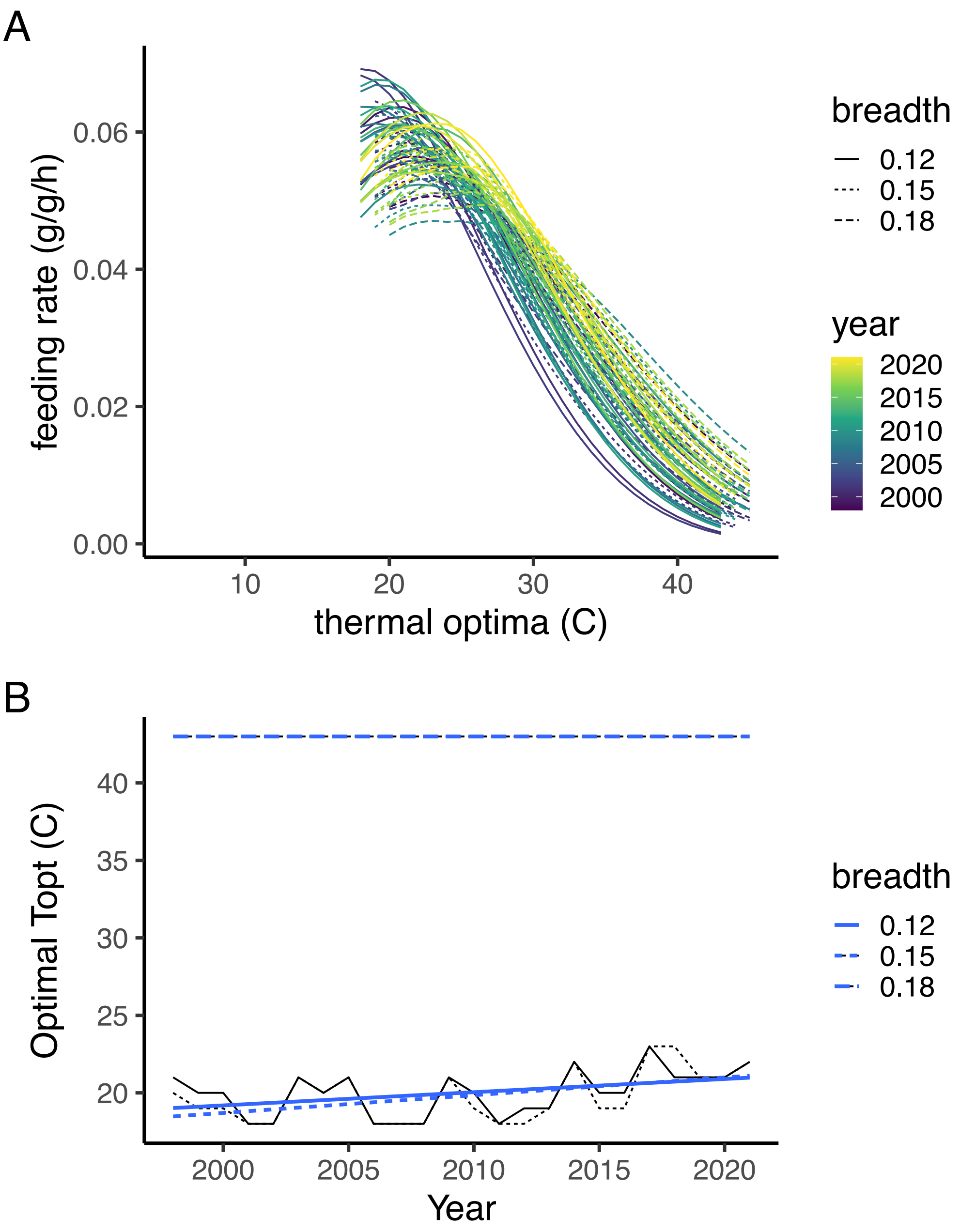


**Figure S5**. (A) We use hourly larval temperatures for Washington *P. rapae* for each year to estimate how thermal optima influences feeding rate and estimate the (B) thermal optima associated with maximum feeding rate. The TPC breath parameter slightly influences (A) the thermal optimal corresponding to maximum feeding. (B) The smaller breadths do not influence the optimal thermal optima across time but for the largest breadth, a high thermal optima increases feeding rate throughout.
